# Opposite roles of Rad5 in DNA damage tolerance: playing in both error-free and mutagenic lesion bypass

**DOI:** 10.1101/2022.01.06.475185

**Authors:** Katarzyna H. Masłowska, Vincent Pagès

## Abstract

DNA Damage Tolerance (DDT) functions to bypass replication-blocking lesions and is divided into two distinct pathways: error-prone Translesion Synthesis (TLS) and error-free Damage Avoidance (DA).

Rad5 is a a multifunctional protein that is involved in these DDT processes. Rad5 contains three well defined domains: a RING domain that promotes PCNA polyubiquitination, a ssDNA-dependent ATPase/helicase domain, and a Rev1-binding domain. Both the RING domain and the ATPase/helicase domain are conserved in human Rad5 ortholog HLTF.

In this study we used domain-specific mutants to address the contribution of each of *Saccharomyces cerevisiae* Rad5 functions to lesion tolerance. Using an assay based on the insertion of a single lesion into a defined locus in the genome of a living yeast cell, we demonstrate that Rad5 plays two opposite roles in lesion tolerance: i) Rad5 favors error-free lesion bypass by activating template switching through polyubiquitination of PCNA; ii) Rad5 is required for TLS by recruiting the TLS polymerase Rev1. We also show that the helicase activity does not play any role in lesion tolerance.

## INTRODUCTION

The DNA of every living cell is constantly threatened by various damaging agents. Despite the efficient action of DNA repair mechanisms, some damage may persist long enough to be present during replication, blocking the replicative polymerases, which threatens genome stability [1]. Therefore, to complete replication, cells need to tolerate the encountered DNA damage. There are two distinct DNA Damage Tolerance (DDT) mechanisms: i) error-prone Translesion Synthesis (TLS), employing specialized low-fidelity DNA polymerases able to insert a few nucleotides opposite the lesion [2]; ii) Damage Avoidance (DA), a generally error-free pathway that relies on homologous recombination (HR) to retrieve the genetic information from the non-damaged sister chromatid [3] (also reviewed in [4,5]).

The balance between error-prone TLS and error-free DA defines the level of mutagenesis during lesion bypass. However, the current understanding of the precise molecular mechanisms regulating the process of DNA Damage Tolerance is far from complete.

In eukaryotes, lesion tolerance is controlled by the ubiquitination of proliferating cell nuclear antigen (PCNA) (reviewed in [6,7]). PCNA monoubiquitination by Rad6 and Rad18 promotes the recruitment of TLS polymerases. Extending this modification to polyubiquitination by Mms2/Ubc13 and Rad5, enables the recombination-mediated mechanisms [8].

Rad5 is a a large multifunctional protein that contains both ubiquitin ligase and ssDNA-dependent ATPase activities [9]. These domains and functions are shared with its human orthologs HLTF and SHPRH [10,11]. As E3 ubiquitin ligase, Rad5 catalyzes PCNA polyubiquitination by bridging PCNA with the E2 (Mms2-Ubc13) and accelerates ubiquitin transfer from the E2 to Ubi-PCNA. It also acts as a bridging factor to bring Ubc13 and Mms2 into contact with the Rad6/Rad18 complex, thereby providing a means to coordinate the distinct ubiquitin-conjugating activities of Rad6 and Ubc13/ Mms2 [12]. As a DNA-dependent ATPase, Rad5 is a member of the DEAD box family of helicases. *In vitro* it has the capacity to catalyze the reversal of replication fork-like structures [13–15], although in yeast fork reversal has been consistently interpreted as a pathological transaction at replication forks that have lost their replication capacity [16]. It has been suggested that Rad5 ATPase activity is important rather for DSB repair [17]. It has been demonstrated that the Rad5 ATPase/helicase activity is not required for PCNA polyubiquitination [18]. Incidentally, the relevant catalytic domains of Rad5 overlap. The RING E3 ligase domain responsible for E2 interaction resides within the helicase domain, inserted between the conserved helicase motifs III and IV [12,19] (Figure 1B).

**Figure 1.**
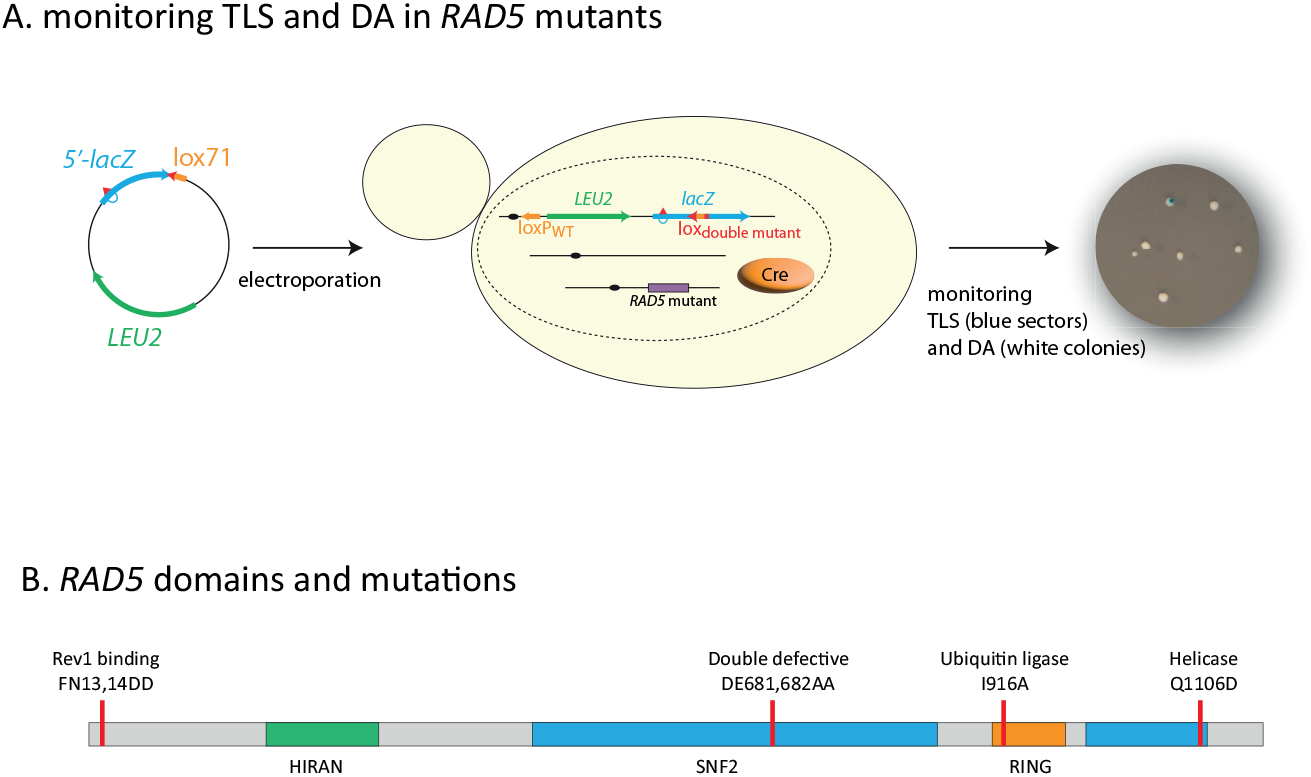
A: outline of the integration system: A non-replicative plasmid containing a single lesion is integrated into one of the yeast chromosomes using Cre/lox site-specific recombination. The integrative vector carrying a selection marker (LEU2) and the 5′-end of the lacZ reporter gene containing a single lesion is introduced into a specific locus of the chromosome with the 3′-end of lacZ. The precise integration of the plasmid DNA into the chromosome restores a functional lacZ gene, enabling the phenotypical detection of TLS and DA events (as blue and white colonies on X-gal indicator media). B: Schematic of RAD5 gene highlighting the helicase domain as well as the RING ubiquitin E3 domain. Mutations used it this study are indicated. Mutation FN13,14DD affects the Rev1 binding site. Mutation DE681,682AA inactivates both the helicase and ubiquitin ligase activity. Mutation I916A inactivates the ubiquitin ligase activity. Mutation Q1106D inactivates the helicase activity.

Rad5 also plays a structural role in the recruitment of TLS polymerases through physical interaction with Rev1 via its N-terminus [20]. The fact that Rad5 is involved in both branches of DDT implies that it may play a role in the pathway choice and balance within DDT. However, we do not know precisely how the three domains of Rad5 (ubiquitin ligase, Rev1-binding, and helicase) affect the choice of the DNA Damage Tolerance pathway.

In this study we used domain-specific mutants to address the contribution of each of the Rad5 domains to the lesion tolerance. We demonstrate that the two critical functions of Rad5 during DNA damage tolerance are the activation of template switching through polyubiquitination of PCNA and the recruitment of TLS. We also show that the helicase activity does not play any role in lesion tolerance.

## RESULTS & DISCUSSION

### Rad5 is involved in Damage Avoidance through its ubiquitin ligase domain

Our group has recently developed an assay based on the insertion of a single lesion into a specific locus in the genome of a living yeast cell, which allows a phenotypical detection of TLS and DA events (as blue and white colonies on X-gal indicator media) (Figure 1A) [21]. In the present work, we have used this assay to determine the role played by the three domains of Rad5 in the balance between TLS and DA.

Using this method, we have introduced a (6-4)TT photoproduct lesion (thymine-thymine pyrimidine (6-4) pyrimidone photoproduct), or a N2dG-AAF (N2-dG-Acetylaminofluorene) adduct in the genome of cells carrying mutations affecting different domains of the Rad5 protein (Figure 1B): i) an allele simultaneously deficient in Ubc13-binding and ATPase/helicase activity (DE681,682AA) named *RAD5DE*^*ubi-helic*^ [13]; ii) the Ubc13-binding RING domain (I916A) named *RAD5IA*^*ubi*^ [19]; iii) the helicase domain (Q1106D) named *RAD5QD*^*helic*^ [18]; iv) and the Rev1-binding domain (FN13,14AA) named *RAD5FN*^*Rev1*^ [20]; v) and a complete deletion of *rad5* gene. The results were compared to the parental strain expressing wild-type *RAD5* gene. In all strains we inactivated *rad14* to avoid repair of the lesion and focus on lesion tolerance mechanisms, and *msh2* to avoid repair of the strand marker that allows to distinguish TLS from DA events.

The mutation affecting Ubc13-binding (*RAD5IA*^*ubi*^) led to a strong increase in TLS at both (6-4)TT photoproduct and N2dG-AAF lesions (Figure 2A and B). This increase in TLS is similar to the one previously observed in the absence of *ubc13* [21,22]. As shown in Figure 4, *RAD5IA*^*ubi*^ shows a high sensitivity to a more global genotoxic stress such as 4NQO (4-Nitroquinoline-1-oxide) treatment, similar to the *ubc13Δ* strain.

**Figure 2:**
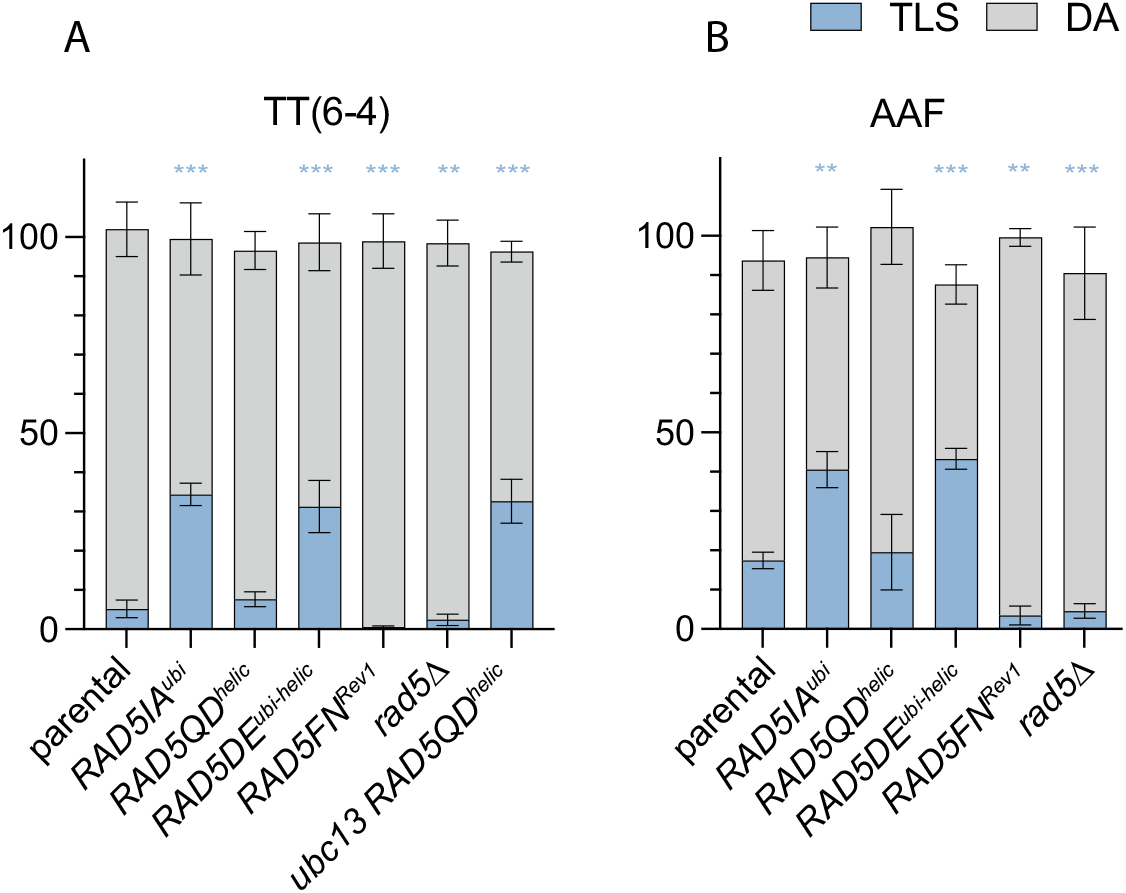
Partitioning of DDT pathways through (6-4)TT and N2dG-AAF lesions in domain-specific Rad5 mutants. Tolerance events represent the percentage of cells able to survive in presence of the integrated lesion compared to the lesion-free control. The data represent the average and standard deviation of at least three independent experiments. Unpaired t-test was performed to compare TLS values from the different mutants to the parental strain (*p < 0.05; **p < 0.005; ***p < 0.0005).

We have previously described a competition between TLS and DA: in the absence of polyubiquitination of PCNA, DA is inhibited favoring TLS. We can note here that the loss of DA in the absence of PCNA polyubiquitination is only partially compensated by an increase in TLS. There is still a proportion of cells surviving using a recombination pathway that is independent of PCNA ubiquitination and that has previously been described as the salvage recombination pathway [23].

### Rad5 helicase function is not involved in damage tolerance

The mutation affecting solely the ATPase/helicase domain (Q1106D) of Rad5 (*RAD5QD*^*helic*^) did not show any effect on the level of TLS and DA at the (6-4)TT photoproduct and N2dG-AAF lesions (Figure 2 A and B). It seems therefore that this function is not involved in the bypass of the tested lesions. One could wonder if the helicase function could act as a backup in the absence of DA. The levels of TLS and DA are the same in the mutant deficient for both helicase and ubiquitin ligase (*RAD5DE*^*ubi-helic*^) as for the mutant deficient solely for the ubiquitin ligase (*RAD5IA*^*ubi*^). Similarly, there is no difference between *ubc13Δ* and *ubc13Δ RAD5QD*^*helic*^ strains. These observations indicate that even in the absence of genuine DA that is dependent on PCNA polyubiquitination (ubc13Δ or *RAD5IA*^*ubi*^), the helicase has no function in damage tolerance. These results are compatible with previous observations from Gallo *et al*. [24] who showed that the helicase mutant had no effect on mutagenesis or survival to HU treatment. The absence of sensitivity of the *RAD5QD*^*helic*^ mutant to 4NQO treatment (Figure 4) confirms that the helicase domain in not involved in lesion tolerance. Previous reports from Chen *et al*. [17] have shown the involvement of the helicase function of Rad5 in double-strand break repair, a role that is independent from its ubiquitin ligase function. *In vitro* experiments have shown the involvement of the helicase domain in fork regression [13], a structure that could favor error-free lesion bypass. *In vitro* experiments have also suggested that Rad5 can facilitate strand invasion-dependent mechanisms in addition to fork regression for the template switching in a Rad51-independent [25]. It appears from our *in vivo* data and others [17,24] that this is not a major pathway *in vivo*, at least for the tested lesions.

It is important to note that previous studies have considered the DE mutant as an ATPase defective strain and evidenced a role for the helicase function in DDT [26]. However, it has since been showed that the DE mutation affects both the helicase and ubiquitin ligase domains [27], therefore the sensitivity that was observed for this mutant was most likely due to the polyubiquitination defect and not the helicase defect.

### Rad5 interaction with Rev1 is required for Polζ-TLS

As we have shown previously, TLS bypass of the (6-4)TT photoproduct relies almost exclusively on the TLS polymerases Rev1 and Pol ζ [21,22]. The bypass of N2dG-AAF lesion is mostly dependent on Rev1 and Pol ζ, while a small part can be performed by pol h [22]. The *RAD5* allele unable to bind Rev1 (*RAD5FN*^*Rev1*^) causes a severe decrease in the level of TLS at both (6-4)TT photoproduct and N2dG-AAF lesions (Figure 2A and B). This indicates that the interaction of Rev1 with Rad5 is critical for its TLS activity *in vivo*. This is in agreement with previous studies which had demonstrated that lack of the Rad5 N-terminal activity severely compromises spontaneous and DNA-damage-induced mutagenesis [20,24]. For the (6-4)TT photoproduct, we confirmed that in the absence of PCNA polyubiquitination in mutants *RAD5DE*^*ubi-helic*^ and *RAD5IA*^*ubi*^, the strong increase in TLS (to a level >30%) was still exclusively due to Rev1-Pol ζ: as observed in Figure 3, the inactivation of *rev1* in the *ubc13*Δ, *RAD5IA*^*ubi*^, or *RAD5DE*^*ubi-helic*^ mutants completely abolishes TLS (≤0.1%). Similarly, inactivation of Rev3 in the *RAD5IA*^*ubi*^ also abolishes TLS.

**Figure 3:**
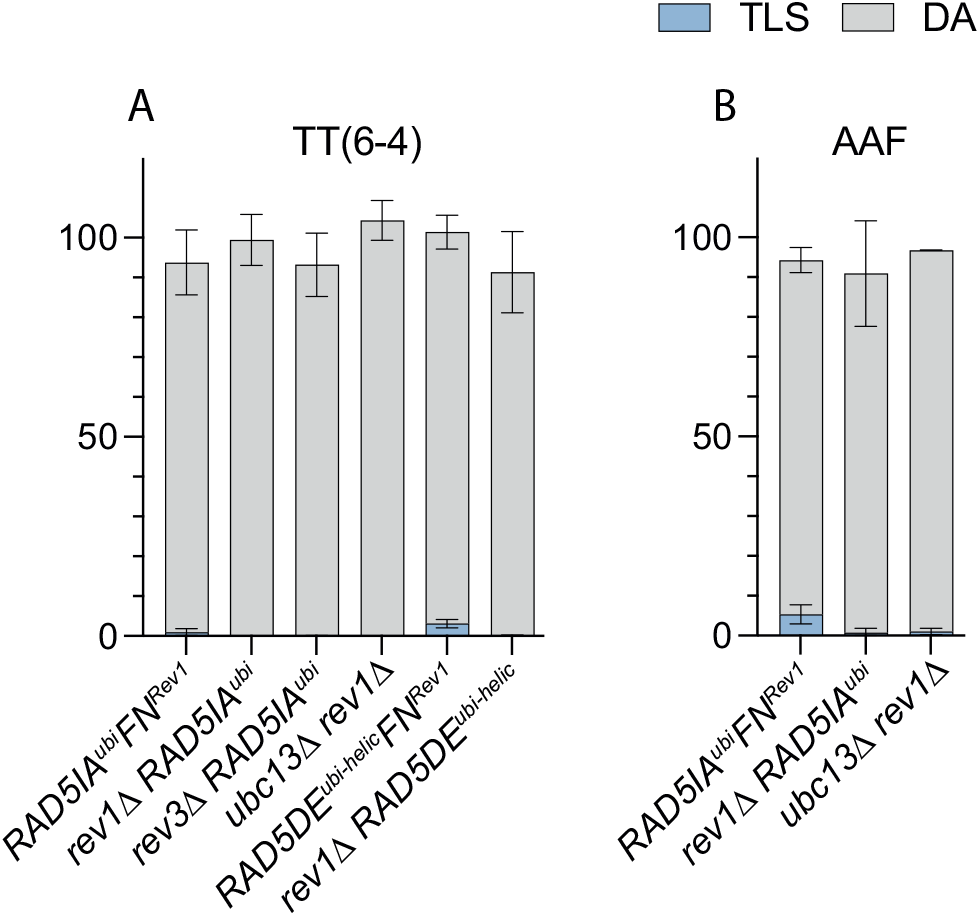
Role of Rev1 and Rad5-Rev1 interaction in the partitioning of DDT pathways through (6-4)TT and N2dG-AAF lesions. Tolerance events represent the percentage of cells able to survive in presence of the integrated lesion compared to the lesion-free control. The data represent the average and standard deviation of at least three independent experiments.

**Figure 4:**
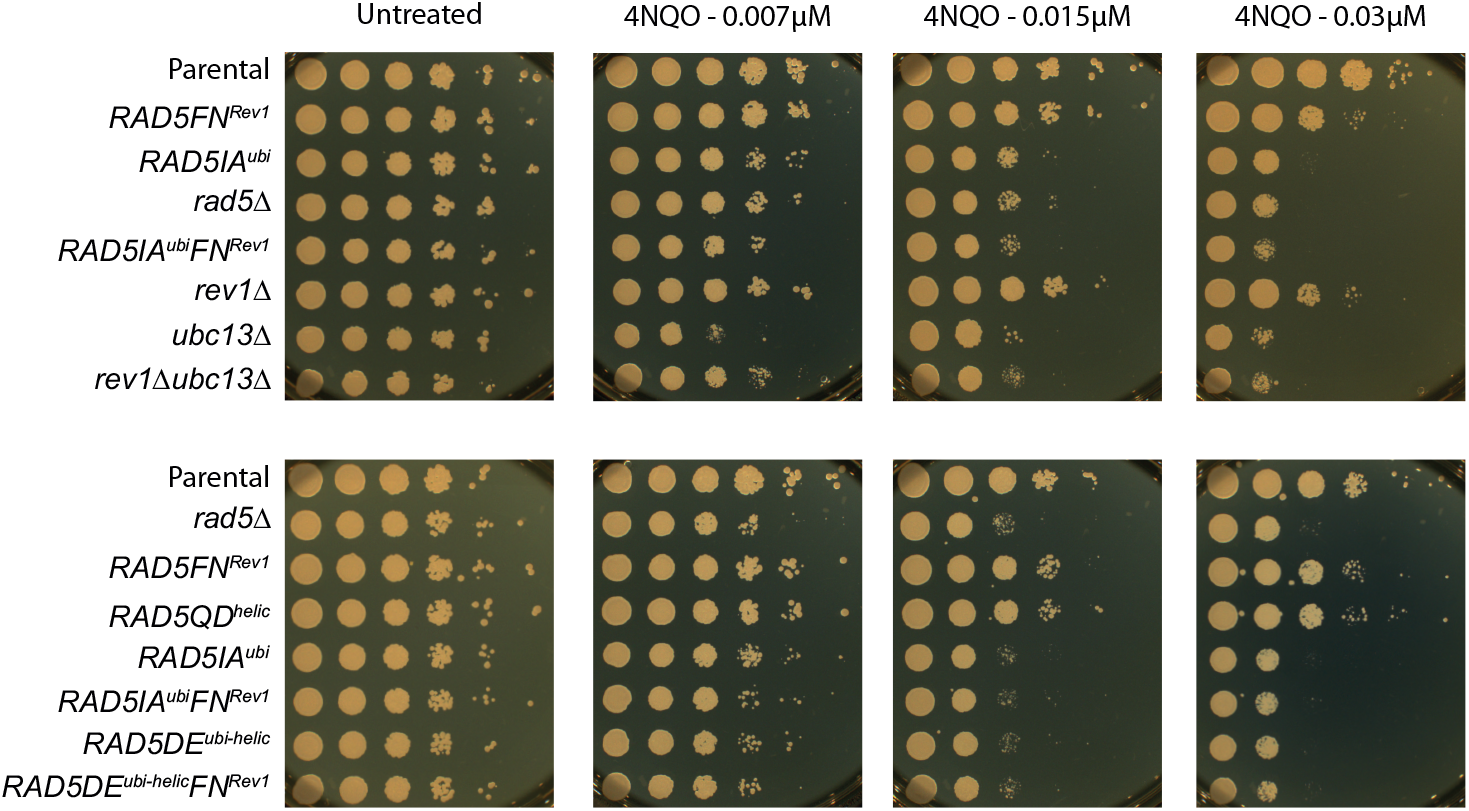
Sensitivity to UV and 4NQO treatment of the different *RAD5* mutants.

When preventing the recruitment of Rev1 by Rad5 (*RAD5FN*^*Rev1*^) in the same mutants *RAD5IA*^*ubi*^ and *RAD5DE*^*ubi-helic*^, we also observed a strong decrease of TLS. However, unlike in the *rev1Δ* strains, TLS is not completely abolished when combining default of PCNA polyubiquitination with *RAD5FN*^*Rev1*^. It appears that when DA is inhibited, Rev1/Pol ζ can access the stalled fork and some TLS can occur despite the absence of recruitment of Rev1 by Rad5.

In response to 4NQO treatment, the *RAD5FN*^*Rev1*^ show the same increased sensitivity as the *rev1Δ* mutant, confirming the importance of Rad5 in recruiting Rev1. It remains however less sensitive than the mutant affecting PCNA polyubiquitination, reflecting the lesser role of TLS for survival over DA. When DA is abolished in the absence of PCNA polyubiquitination (strains *ubc13Δ, RAD5IA*^*ubi*^ or *RAD5DE*^*ubi-helic*^), the further inactivation of rev1 either directly (*rev1Δ* strain) or through the lack of recruitment by Rad 5 (*RAD5FN*^*Rev1*^) does not further increase the sensitivity to 4NQO, indicating again the minor role of TLS for survival.

While the complete deletion of *rad5* leads to an increased sensitivity to 4NQO (Figure 4), it does not lead to a drastic phenotype when monitoring the bypass of a single (6-4)TT photoproduct or N2dG-AAF lesions compared to WT *RAD5* (Figure 2A and B). We observed a significant decrease in TLS for the N2dG-AAF lesions, and a very moderate decrease for the TT(6-4) photoproduct compared to the parental strain. It is important to note that in the *rad5*Δ strain, no polyubiquitination of PCNA occurs: we could therefore expect in these strains a strong increase of TLS as observed in the *ubc13*Δ, *RAD5IA*^*ubi*^, or *RAD5DE*^*ubi-helic*^ mutants [21,22]. However, due to the absence of Rad5 and its function of recruiting Rev1, TLS does not increase in this strain. Overall, the loss of DA due to the absence of PCNA-ubiquitination could not be compensated by an increase in TLS in the absence of Rev1 recruitment, and is therefore compensated by an increase in the salvage recombination pathway.

## CONCLUSION

From these data, we can draw the following model: Rad5 is recruited to the replication fork through its interaction with both Rad18 and PCNA [12], where it recruits Rev1 to allow TLS [20] and polyubiquitinates PCNA to allow DA. If Rad5 is unable to polyubiquitinate PCNA (*RAD5DE*^*ubi-helic*^ or *RAD5IA*^*ubi*^ mutant), it will recruit Rev1 and permits a high level of TLS as DA is inhibited. If Rad5 is unable to interact with Rev1 (*RAD5FN*^*Rev1*^ mutant), then Rev1-Polζ-TLS will be strongly reduced. A Rad5 protein defective for its two main functions (*RAD5IA*^*ubi*^, *FN* ^*Rev1*^) has a phenotype similar to a complete deletion of Rad5: it shows the same level of TLS and DA at the (6-4)TT photoproduct (compare Figure 2B and Figure 3), and similar sensitivity to 4NQO (Figure 4). In the absence of Rad5, Rev1 can access the replication fork and perform TLS, but with a lower efficiency as it is not actively recruited by Rad5. In conclusion, we have shown that Rad5 plays two critical and opposite roles in lesion tolerance: i) through its ubiquitin ligase activity, Rad5 promotes error-free lesion bypass by damage avoidance, and ii) through its interaction with Rev1, it promotes Rev1/Pol ζ -dependent error-prone TLS. The helicase activity that has been suggested to favor error-free bypass by promoting fork regression does not seem to play a key role in the tolerance of isolated lesions [17].

## MATERIAL AND METHODS

### Strains and media

All strains used in the present study are derivative of strain EMY74.7 [28] (MATa his3-Δ1 leu2-3,112 trp1-Δ ura3-Δ met25-Δ phr1-Δ rad14-Δ msh2Δ::hisG). In order to study tolerance events, all strains are deficient in repair mechanisms: nucleotide excision repair (*rad14*), photolyase (*phr1*), and mismatch repair system (*msh2*). Gene disruptions were achieved using PCR-mediated seamless gene deletion [29] or URAblaster [30] techniques. Rad5 point mutations were created using the delitto perfetto method [31]. All strains used in the study are listed in Table 1.

**Table 1:**
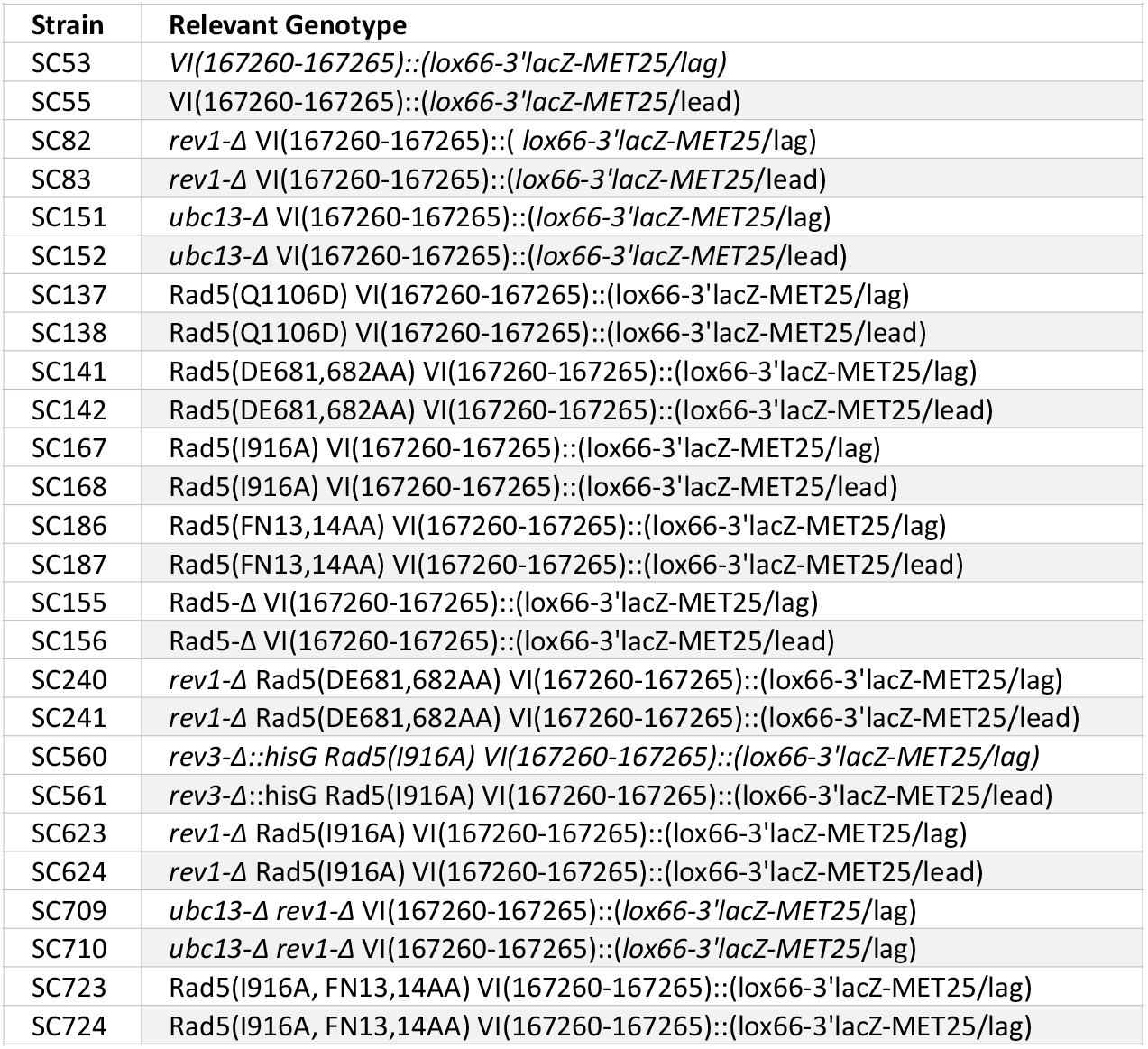
Strains used in the study. All strains are: MATa his3-Δ1 leu2-3,112 trp1-Δ ura3-Δ met25-Δ rad14-Δ phr1-Δ msh2Δ::hisG.

### Integration system

Integration of plasmids carrying 6-4 (TT) / N2dG-AAF lesions (or control plasmids without lesion) and result analysis was performed as previously described [21]. All experiments were performed at least in triplicate. Graphs and statistical analysis were done using GraphPad Prism applying unpaired t-test. Bars represent the mean value ± s.d.

### Spotting assay

Overnight cultures of strains carrying Rad5 point mutations in YPD were adjusted to an OD_600_ value of 1. Volume of 10 μl from 10-fold serial dilutions of OD_600_-adjusted cultures were spotted on YPD agar plates containing different concentrations of 4-NQO (0 μM, 0,015 μM, 0,03 μM).

## Declaration of interests

The authors declare that they have no known competing financial interests or personal relationships that could have appeared to influence the work reported in this paper.

## Aknowledgement

This work as supported by Fondation pour la Recherche Médicale [FRM - EQU2 01903007797]; Fondation de France (to K.M.).

## Author contribution

V.P. and K.M. conceptualization; V.P. and K.M. data curation; V.P. and K.M. formal analysis; K.M. investigation; VP. supervision; V.P. and K.M. funding acquisition; V.P. and K.M. writing-original draft.

## Notes

### Competing Interest Statement

The authors have declared no competing interest.

### Summary of Updates

We have added data combining mutations in ubiquitination defect and Rev1 binding defect

## REFERENCES

1. Friedberg EC, Walker GC, Siede W, Wood RD, Schultz R a., Ellenburger T. DNA repair and mutagenesis. 2nd ed. Washington DC: ASM Press; 2006. doi:10.1097/01.shk.0000232588.61871.ff

2. Sale JE, Lehmann AR, Woodgate R. Y-family DNA polymerases and their role in tolerance of cellular DNA damage. Nature Reviews Molecular Cell Biology. 2012;13: 141–152. doi:10.1038/nrm3289.Y-family

3. Branzei D. Ubiquitin family modifications and template switching. FEBS Letters. 2011;585: 2810–2817. doi:10.1016/j.febslet.2011.04.053

4. Branzei D, Szakal B. DNA damage tolerance by recombination: Molecular pathways and DNA structures. DNA Repair. 2016;44: 68–75. doi:10.1016/j.dnarep.2016.05.008

5. Waters LS, Minesinger BK, Wiltrout ME, D’Souza S, Woodruff R V., Walker GC. Eukaryotic Translesion Polymerases and Their Roles and Regulation in DNA Damage Tolerance. Microbiology and Molecular Biology Reviews. 2009;73: 134–154. doi:10.1128/MMBR.00034-08

6. Andersen PL, Xu F, Xiao W. Eukaryotic DNA damage tolerance and translesion synthesis through covalent modifications of PCNA. Cell Research. 2008;18: 162– 173. doi:10.1038/cr.2007.114

7. Che J, Hong X. PCNA Ubiquitylation : Instructive or Permissive to DNA Damage Tolerance Pathways? 2021;13: 2–7.

8. Hoege C, Pfander B, Moldovan G-L, Pyrowolakis G, Jentsch S. RAD6-dependent DNA repair is linked to modification of PCNA by ubiquitin and SUMO. Nature. 2002;419: 135–141. doi:10.1038/nature00991

9. Unk I, Hajdú I, Blastyák A, Haracska L. Role of yeast Rad5 and its human orthologs, HLTF and SHPRH in DNA damage tolerance. DNA Repair. 2010;9: 257–267. doi:10.1016/j.dnarep.2009.12.013

10. Unk I, Hajdú I, Fátyol K, Szakál B, Blastyák A, Bermudez V, et al. Human SHPRH is a ubiquitin ligase for Mms2-Ubc13-dependent polyubiquitylation of proliferating cell nuclear antigen. Proc Natl Acad Sci U S A. 2006;103: 18107–18112. doi:10.1073/pnas.0608595103

11. Unk I, Hajdu I, Fatyol K, Hurwitz J, Yoon J-H, Prakash S, et al. Human HLTF functions as a ubiquitin ligase for proliferating cell nuclear antigen polyubiquitination. Proceedings of the National Academy of Sciences. 2008;105: 3768–3773. doi:10.1073/pnas.0800563105

12. Ulrich HD, Jentsch S. Two RING finger proteins mediate cooperation between ubiquitin-conjugating enzymes in DNA repair. EMBO J. 2000;19: 3388–97. doi:10.1093/emboj/19.13.3388

13. Blastyák A, Pintér L, Unk I, Prakash L, Prakash S, Haracska L. Yeast Rad5 Protein Required for Postreplication Repair Has a DNA Helicase Activity Specific for Replication Fork Regression. Molecular Cell. 2007;28: 167–175. doi:10.1016/j.molcel.2007.07.030

14. Parker JL, Ulrich HD. Mechanistic analysis of PCNA poly-ubiquitylation by the ubiquitin protein ligases Rad18 and Rad5. EMBO Journal. 2009;28: 3657–3666. doi:10.1038/emboj.2009.303

15. Carlile CM, Pickart CM, Matunis MJ, Cohen RE. Synthesis of free and proliferating cell nuclear antigen-bound polyubiquitin chains by the RING E3 ubiquitin ligase Rad5. Journal of Biological Chemistry. 2009;284: 29326–29334. doi:10.1074/jbc.M109.043885

16. Neelsen KJ, Lopes M. Replication fork reversal in eukaryotes: from dead end to dynamic response. Nat Rev Mol Cell Biol. 2015;16: 207–220. doi:10.1038/nrm3935

17. Chen S, Davies A, Sagan D, Ulrich HD. The RING finger ATPase Rad5p of Saccharomyces cerevisiae contributes to DNA double-strand break repair in a ubiquitin-independent manner. Nucleic Acids Research. 2005;33: 5878–5886. doi:10.1093/nar/gki902

18. Choi K, Batke S, Szakal B, Lowther J, Hao F, Sarangi P, et al. Concerted and differential actions of two enzymatic domains underlie Rad5 contributions to DNA damage tolerance. Nucleic Acids Research. 2015;43: 2666–2677. doi:10.1093/nar/gkv004

19. Ulrich HD. Protein-protein interactions within an E2-RING finger complex: Implications for ubiquitin-dependent DNA damage repair. Journal of Biological Chemistry. 2003;278: 7051–7058. doi:10.1074/jbc.M212195200

20. Xu X, Lin A, Zhou C, Blackwell SR, Zhang Y, Wang Z, et al. Involvement of budding yeast Rad5 in translesion DNA synthesis through physical interaction with Rev1. Nucleic Acids Research. 2016;44: 5231–5245. doi:10.1093/nar/gkw183

21. Maslowska KH, Laureti L, Pagès V. iDamage: a method to integrate modified DNA into the yeast genome. Nucleic Acids Res. 2019;47: e124. doi:10.1093/nar/gkz723

22. Masłowska KH, Villafañez F, Laureti L, Iwai S, Pagès V. Eukaryotic stress induced mutagenesis is limited by a local control of Translesion Synthesis. Nucleic Acids Research. 2022. Available: https://doi.org/10.1093/nar/gkac044

23. Pfander B, Moldovan G-LL, Sacher M, Hoege C, Jentsch S, Sacher M, et al. SUMO-modified PCNA recruits Srs2 to prevent recombination during S phase. Nature. 2005;436: 428–433. doi:10.1038/nature03665

24. Gallo D, Kim T, Szakal B, Saayman X, Narula A, Park Y, et al. Rad5 Recruits Error-Prone DNA Polymerases for Mutagenic Repair of ssDNA Gaps on Undamaged Templates. Molecular Cell. 2019;73: 1–15. doi:10.1016/j.molcel.2019.01.001

25. Burkovics P, Sebesta M, Balogh D, Haracska L, Krejci L. Strand invasion by HLTF as a mechanism for template switch in fork rescue. Nucleic Acids Research. 2014;42: 1711–1720. doi:10.1093/nar/gkt1040

26. Gangavarapu V, Haracska L, Unk I, Johnson RE, Prakash S, Prakash L. Mms2-Ubc13-dependent and -independent roles of Rad5 ubiquitin ligase in postreplication repair and translesion DNA synthesis in Saccharomyces cerevisiae. Mol Cell Biol. 2006;26: 7783– 90. doi:10.1128/MCB.01260-06

27. Ball LG, Xu X, Blackwell SR, Hanna MD, Lambrecht AD, Xiao W. The Rad5 helicase activity is dispensable for error-free DNA post-replication repair. DNA Repair. 2014;16: 74–83. doi:10.1016/j.dnarep.2014.02.016

28. Johnson RE, Torres-Ramos CA, Izumi T, Mitra S, Prakash S, Prakash L. Identification of APN2, the Saccharomyces cerevisiae homolog of the major human AP endonuclease HAP1, and its role in the repair of abasic sites. Genes and Development. 1998;12: 3137– 3143. doi:10.1101/gad.12.19.3137

29. Akada R, Kitagawa T, Kaneko S, Toyonaga D, Ito S, Kakihara Y, et al. PCR-mediated seamless gene deletion and marker recycling in Saccharomyces cerevisiae. Yeast. 2006;23: 399–405. doi:10.1002/yea.1365

30. Alani E, Cao L, Kleckner N. A method for gene disruption that allows repeated use of URA3 selection in the construction of multiply disrupted yeast strains. Genetics. 1987;116: 541–545. doi:10.1534/genetics.112.541.test

31. Storici F, Resnick MA. The Delitto Perfetto Approach to In Vivo Site-Directed Mutagenesis and Chromosome Rearrangements with Synthetic Oligonucleotides in Yeast. Methods in Enzymology. 2006;409: 329–345. doi:10.1016/S0076-6879(05)09019-1

